# Inhibitory effect of Ephedra herba on human norovirus infection in human intestinal organoids

**DOI:** 10.1101/2023.03.08.531477

**Authors:** Tsuyoshi Hayashi, Kosuke Murakami, Hirokazu Ando, Sayuri Ueno, Sakura Kobayashi, Masamichi Muramatsu, Takashi Tanikawa, Masashi Kitamura

**Author notes:** Correspondence to: Tsuyoshi Hayashi, Takashi Tanikawa, Masashi Kitamura.

## Abstract

Human norovirus (HuNoV) is a major cause of acute gastroenteritis and foodborne diseases worldwide with public health concern, yet no antiviral therapies have been developed. In this study, we aimed to screen crude drugs, which are components of Japanese traditional medicine, ‘‘Kampo” to see their effects on HuNoV infection using a reproducible HuNoV cultivation system, stem-cell derived human intestinal organoids/enteroids (HIOs). Among the 23 crude drugs tested, Ephedra herba significantly inhibited HuNoV infection in HIOs. A time-of-drug addition experiment indicated that this crude drug likely targets post-entry step for the inhibition. To our knowledge, this is the first anti-HuNoV inhibitor screen targeting crude drugs, and Ephedra herba was identified as a novel inhibitor candidate that merits further study.

## 1. Introduction

Human norovirus (HuNoV) is a highly contagious pathogen that causes acute gastroenteritis and foodborne diseases globally, and thereby poses a threat to public health. Despite its clinical importance, no approved antiviral therapies or vaccines are currently available. Among the 10 genogroups that contain 49 genotypes as classified by the Norovirus Classification Working Group [1], GII.4 viruses (genogroup II, genotype 4) are major causes of norovirus outbreak in human populations worldwide [2].

Since its discovery in 1970s, HuNoV basic research using a live virus remained relatively unexplored until recently, because of the longstanding lack of *in vitro* cultivation system. Recently, several laboratory HuNoV cultivation systems have been established [3–5]. A stem-cell derived human intestinal organoid/enteroid (HIO) is one model that supports HuNoV replication reproducibly *in vitro* and is being used to investigate the molecular mechanisms of viral infection [5].

Kampo medicine, a Japanese herbal medicine adapted from traditional Chinese medicine, is being prescribed for patients with various diseases and is supported by the insurance system in Japan [6,7]. Kampo formulations consist a mixture of ‘‘crude drugs” extracted from natural products such as plant components (e.g., leaves, flowers, and branches), minerals, or insects. For example, one of the well-known Kampo, called maoto consists of a mixture of four crude drugs/herbs: Ephedrae Herba, Cinnamomi Cortex, Armeniacae Semen, and Glycyrrhize Radix.

Several crude drugs including Ephedrae Herba have been reported to exhibit inhibitory effects against particular viruses such as SARS-CoV-2, influenza virus, and respiratory syncytial virus (RSV) [8–10]. However, to our knowledge, the effects of crude drugs on HuNoV infection have not yet been tested. In this study, we aimed to screen these crude drugs to determine their effects on HuNoV infection using the HIO culture system.

## 2. Materials and methods

### 2-1. Crude drugs

In total, 23 crude drugs (Table S1) were tested for their effects against HuNoV infection. Each crude drug (10 g) was refluxed with 300 mL of 70% EtOH for 1 hr, and the resultant extracts were dried by evaporation as described previously [11]. The samples were then dissolved in dimethyl sulfoxide (DMSO) to a final concentration of 10 mg/mL and were kept at −30 or −80 °C before use.

### 2-2. HIO culture

A jejunal HIO J2 line, obtained from Baylor College of Medicine under a material transfer agreement, was embedded with Matrigel and cultured as 3-dimensional (3D) HIO in complete medium with growth factors [CMGF(+)] or IntestiCult Organoid Growth Medium (STEMCELL) as described previously [5,12,13]. To prepare differentiated, 2-dimensional (2D) HIO monolayers for HuNoV infection experiments, 3D HIOs (passages 21–29) were dissociated with TrypLE Express (Thermo Fisher) and seeded onto collagen IV-coated 96-well plates (approximately 10^5^ cells/well) in IntestiCult Organoid Growth Medium supplemented with ROCK inhibitor Y-27632 (10 μM; Sigma) for 2-3 days. The medium was then replaced with IntestiCult Organoid Differentiation Medium (STEMCELL)’ and the cells were cultured for another 2 days [5,12,13].

### 2-3. HuNoV infection using HIO culture system

The 2D HIO monolayers in 96-well plate were inoculated with 10% stool filtrate containing 4.3 x 10^5^ genome equivalents (GEs) of GII.3 [GII.P21] (TCH04-577) or 5.7 x 10^5^ GEs of GII.4 [GII.P16] HuNoV in the presence of DMSO (non-drug control) or crude drugs at the indicated concentrations for 1 hr. The cells were then washed twice with complete medium without growth factors [CMGF(-)], and cultured in IntestiCult Organoid Differentiation Medium in the presence of DMSO or crude drugs until 24 hrs post-infection (hpi). Noted that the 500 μM GCDCA, which is indispensable for GII.3 HuNoV infection and enhances GII.4 HuNoV infection [5,14], was added to the medium throughout the duration of the experiment.

A time of drug addition experiment was performed as described previously [15]. Briefly, the 2D HIO monolayers were infected with GII.4 HuNoV-containing 10% stool filtrate under three different conditions (Figure 3A): (a) the cells were treated with the drug throughout the duration of the experiment (whole); (b) the drug was present in the first 3 hrs after infection, which likely reflects viral entry into the cells, but prior to replication; (c) the drug was added to the medium after 3 hpi, which likely reflects post-entry step in which the infecting virus replicates in the cells, and assembles and releases new particles that may re-infect new cells.

### 2-4. Quantification of HuNoV RNA and cytotoxicity assay

After 24 hpi, the cells and medium were harvested and subjected to RNA extraction using the Direct-zol RNA MiniPrep Kit (Zymo Research) or Direct-zol-96 RNA Kit (Zymo Research). The samples were used for quantitating HuNoV RNA GEs by means of reverse transcription-quantitative PCR (RT-qPCR) analysis using a TaqMan Fast Virus 1-Step Master Mix (Thermo Fisher) and GII specific primer/probe sets, as reported previously [5,12,13,16]. The medium was also used for measuring cytotoxicity using Cytotoxicity LDH assay kit-WST (Dojindo) [12].

### 2-5. Statistical analysis

Statistical analysis was performed using two-tailed Student t test or ANOVA followed by Dunnett’s multiple-comparison test using the GraphPad Prism 9 software and *P* values of < 0.05 was considered statistically significant.

## 3. Results

### 3-1. Screening of crude drugs using HIOs

We initially screened 23 crude drugs to determine their effects upon HuNoV infection using the HIO culture system. Two-dimensional, differentiated HIO monolayers were infected with GII.4 HuNoV in the presence of either DMSO as a control or crude drugs (50 μg/ml) for 24 hrs. The cells and/or medium were then harvested to determine viral replication and cytotoxicity as described in Materials and Methods (Fig. 1). As expected, HuNoV replicated in DMSO-treated HIOs showing a 215-fold increase in GEs at 24 hpi as compared to that at 1 hpi. Among the 23 crude drugs tested, Forsythiae Fructus showed the strongest antiviral effect, while this effect was highly likely due to cytotoxicity (Fig. 1). In contrast, Ephedra herba inhibited HuNoV infection at a similar level to Forsythiae Fructus, but without compromising cell viability (Fig. 1).

**Figure 1.**
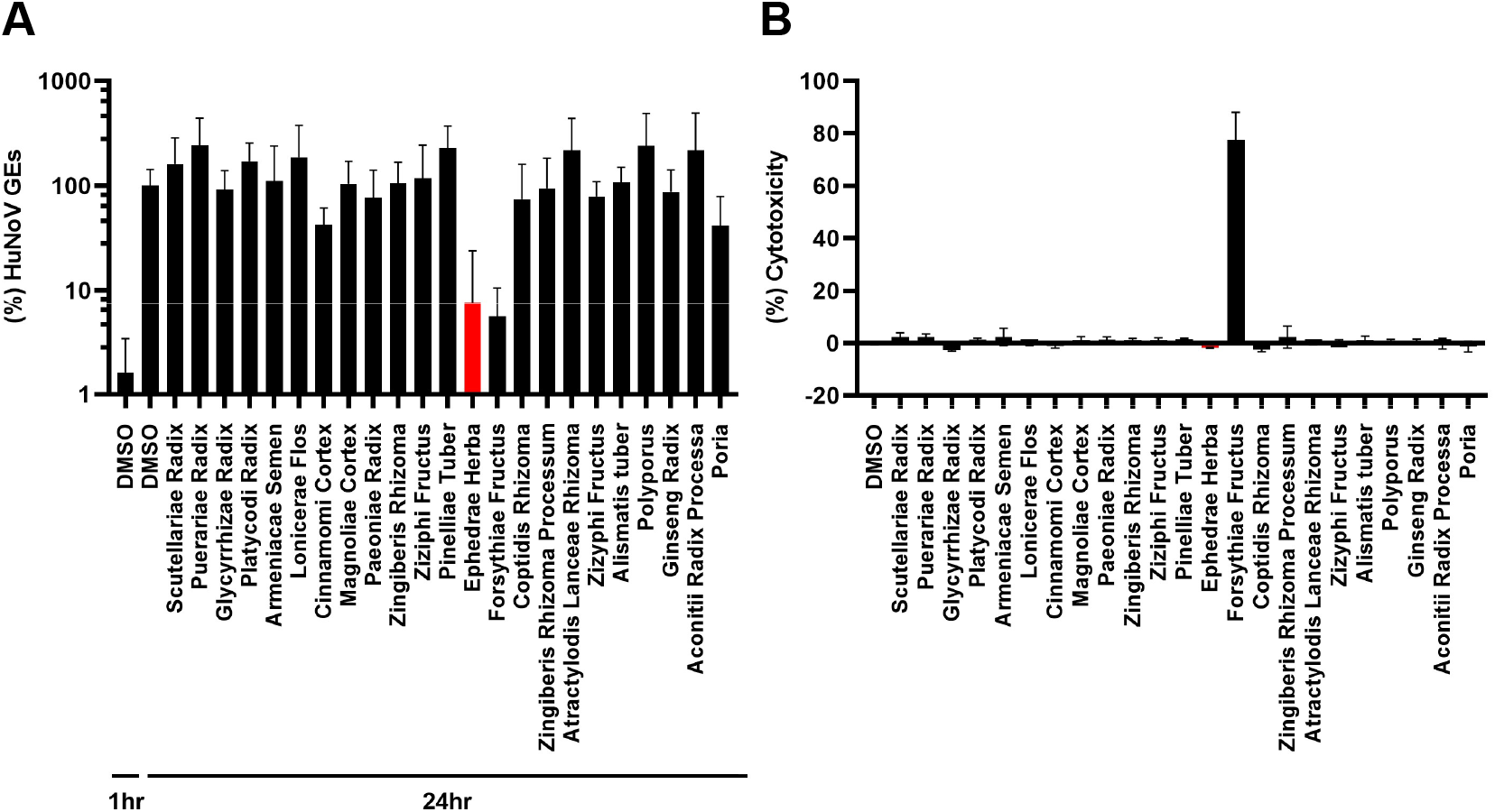
Effect of 23 crude drugs on HuNoV infection in HIOs. Differentiated HIO monolayers were inoculated with GII.4 HuNoV-containing stool filtrate in the presence of the indicated crude drug (50 μg/mL) for 1hr. The cells were washed and further incubated in medium containing crude drug in a 100 μL volume until 24 hpi. The cells and 75 μL of supernatant at 1 hpi or 24 hpi were subjected to RNA extraction followed by RT-qPCR analysis to determine viral GEs (A). The 25μL of supernatant at 24 hpi was used to measure cytotoxicity using lactate dehydrogenase assay (B). The experiments were performed at least twice with more than two technical replicates. The results were normalized to the DMSO control at 24 hpi and are presented as the mean ± standard deviation (n ≥ 4).

### 3-2. Inhibitory effect of Ephedra herba on HuNoV infection

Next, we performed a titration experiment to further validate the effect of Ephedra herba on HuNoV infection. We found a dose-dependent inhibition of GII.4 HuNoV replication compared to DMSO control with no cytotoxicity (Fig. 2A and B). A similar inhibitory effect was seen for the replication of GII.3 HuNoV (Fig. 2C and D).

**Figure 2.**
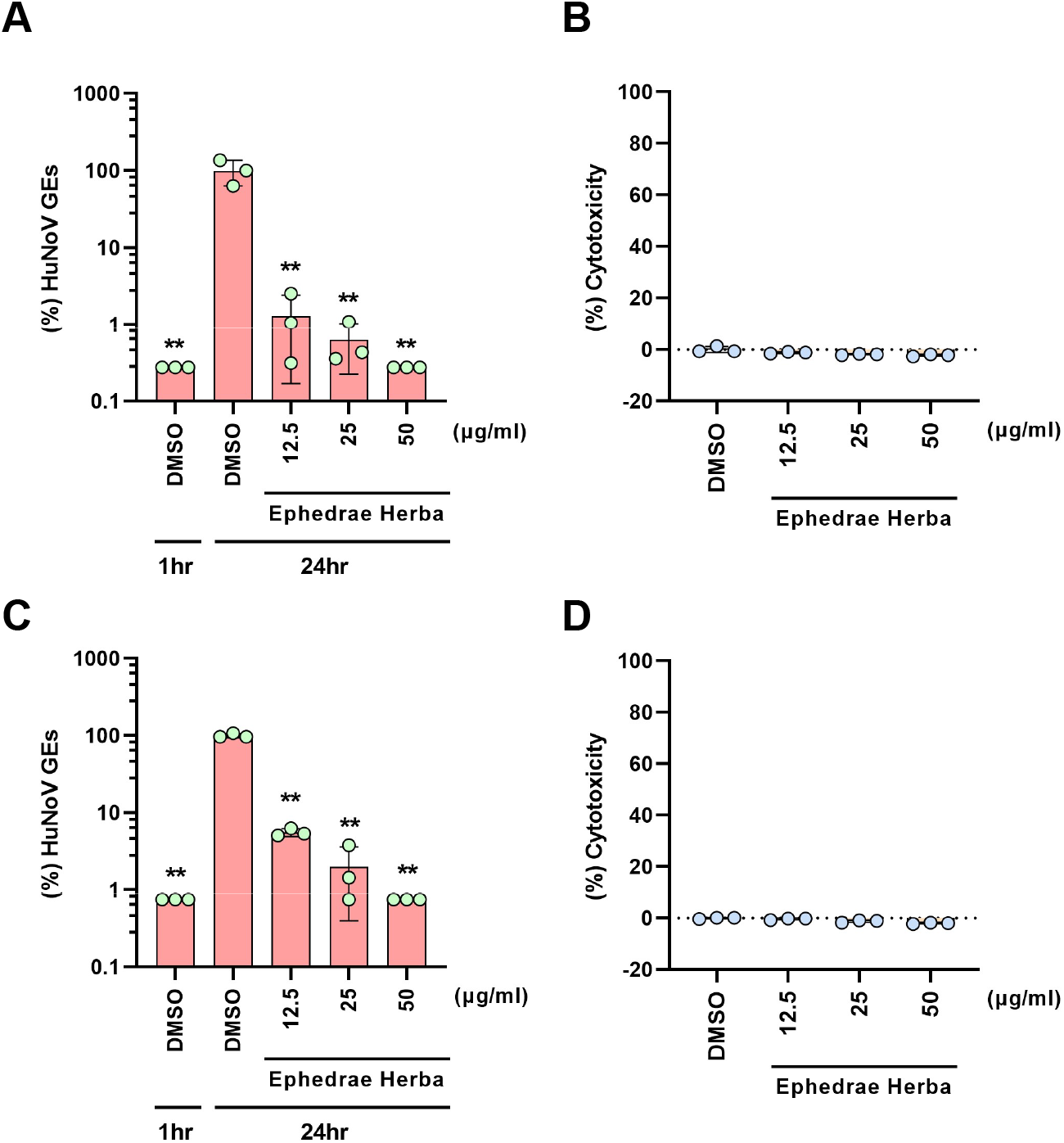
Ephedra herba inhibits GII.3 or GII.4 HuNoV infection in HIOs. The HIO monolayers were inoculated with 10% stool filtrate containing GII.3 (A and B) or GII.4 HuNoV (C and D) in the presence of Ephedra herba at the indicated concentrations and were cultured until 24 hpi. The HuNoV GEs (A and C) and cytotoxicity (B and D) were determined as in Fig. 1 and were normalized to the DMSO control. The experiments were performed twice with three technical replicates, and the results are presented as mean ± standard deviation, with data from one representative experiment (n = 3). ** *p* < 0.01, two-tailed Student’s t-test.

We then tried to determine if Ephedra herba targets HuNoV’s entry or post-entry step by comparing viral replication under three different treatment conditions as shown in Fig. 3A. We used 2’-C-Methylcytidine (2-CMC) as a control, which has been recently reported to inhibit HuNoV replication in HIO [12] and is expected to target a post-entry step, as this compound is a nucleoside polymerase inhibitor against various viruses such as hepatitis C virus and murine norovirus. As expected, 2-CMC completely inhibited viral replication in HIOs under the conditions in which the drug was present throughout the infection period (a) and between 3 and 24 hrs (c), but not in the first 3 hrs after the infection (b) (Fig. 3B). The Ephedra herba significantly inhibited viral replication in HIOs under all conditions tested, while its inhibitory effect in (c) was much stronger than that in (b), suggesting that it preferentially targets the post-entry step for the inhibition (Fig. 3B).

**Figure 3.**
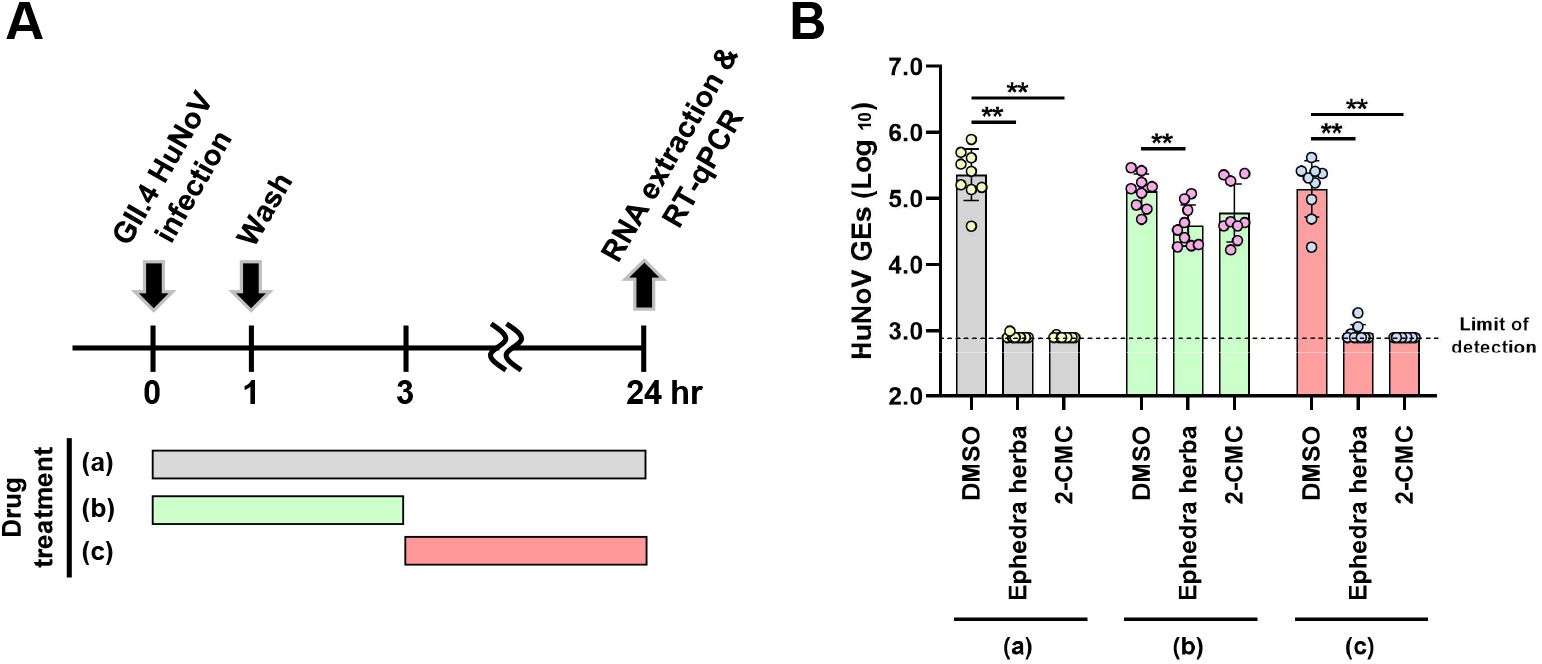
Time-of-drug addition assay. The HIO monolayers were inoculated with GII.4 HuNoV-containing stool filtrate and were cultured until 24 hpi. The cells were treated with DMSO, Ephedra herba (50 μg/mL), or 2-CMC (389 μM) throughout the duration of the experiment (a), in the first 3 hpi (b), or between 3 hpi and 24 hpi (c). After the cells and media were harvested, the HuNoV GEs were determined as in Fig. 1. The experiments were performed three times with three technical replicates, and the results are presented as the mean ± standard deviation (n = 9). The limit of detection in RT-qPCR analysis is 2.9 log_10_ GEs/well. ** *p* < 0.01 *versus* DMSO control, one-way ANOVA followed by Dunnett’s multiple-comparison test.

## 4. Discussion

As reproducible HuNoV cultivation systems were established recently, studies on anti-HuNoV drug discovery are in their infancy. Till date, only few compounds or extracts of natural products have been reported to inhibit a *de novo* HuNoV infection [12,17–19]. In this study, to our knowledge, we performed the first screening of crude drugs, all of which are components of Kampo medicine, to assess their effects on HuNoV infection and identified Ephedra herba as a novel anti-HuNoV drug candidate.

The mechanism by which Ephedra herba inhibits HuNoV infection remains to be determined in this study. A time-of-drug addition experiment suggested that this crude drug likely targets the post-entry step for the inhibition (Fig. 3). It has been reported that Ephedra herba inhibits influenza replication in culture cells, likely due to the inhibition of endosomal acidification [10], whereas it inhibits RSV infection through binding to its envelope G protein, which results in inhibition of its binding to the RSV’s attachment receptor, CX3CR1 [8]. It should be considered that Ephedra herba contains a wide variety of molecules including ephedrine alkaloids (e.g., ephedrine, pseudoephedrine) and polyphenols (e.g., flavonoids, tannins). This implies that active molecule(s) eliciting antiviral activity in Ephedra herba might vary among viruses. Further studies to pinpoint active molecule(s) that inhibit HuNoV infection are needed to better understand how Ephedra herba and its active molecule(s) work, which will contribute to the development of anti-HuNoV agent.

## Author contributions

T.H., T.T., and M.K. conceived the study aim, designed the experiments and wrote the manuscript. T.H., K.M., S.U., S.K., T.T., and M.K. performed the experiments and analyzed the data. H.A., and M.M. provided resources and advice for the study, discussed the results, and critically reviewed the manuscript. All the authors have approved the final version of the manuscript.

## Acknowledgements

We thank Chizuko Hirano (National Institute of Infectious Diseases, Japan) for technical assistance. We also thank Dr. Mary K. Estes (Baylor College of Medicine, USA) for critically reading and technical supports. We thank Editage (www.editage.com) for editing this manuscript.

---

**Table S1.** Crude drugs used in this study.

| Latin name | Distributor |
| --- | --- |
| Scutellariae Radix | Uchida Wakanyaku Co.,Ltd. (Tokyo, Japan) |
| Puerariae Radix | Uchida Wakanyaku Co.,Ltd. (Tokyo, Japan) |
| Glycyrrhizae Radix | Uchida Wakanyaku Co.,Ltd. (Tokyo, Japan) |
| Platycodi Radix | Uchida Wakanyaku Co.,Ltd. (Tokyo, Japan) |
| Armeniacae Semen | Tochimoto Tenkaido Co. Ltd. (Osaka, Japan) |
| Lonicerae Flos | Tochimoto Tenkaido Co. Ltd. (Osaka, Japan) |
| Cinnamomi Cortex | Uchida Wakanyaku Co.,Ltd. (Tokyo, Japan) |
| Magnoliae Cortex | Tochimoto Tenkaido Co. Ltd. (Osaka, Japan) |
| Paeoniae Radix | Uchida Wakanyaku Co.,Ltd. (Tokyo, Japan) |
| Zingiberis Rhizoma | Uchida Wakanyaku Co.,Ltd. (Tokyo, Japan) |
| Ziziphi Fructus | Uchida Wakanyaku Co.,Ltd. (Tokyo, Japan) |
| Pinelliae Tuber | Tochimoto Tenkaido Co. Ltd. (Osaka, Japan) |
| Ephedrae Herba | Uchida Wakanyaku Co.,Ltd. (Tokyo, Japan) |
| Forsythiae Fructus | Tochimoto Tenkaido Co. Ltd. (Osaka, Japan) |
| Coptidis Rhizoma | Tochimoto Tenkaido Co. Ltd. (Osaka, Japan) |
| Zingiberis Rhizoma Processum | Tochimoto Tenkaido Co. Ltd. (Osaka, Japan) |
| Atractylodis Lanceae Rhizoma | Tochimoto Tenkaido Co. Ltd. (Osaka, Japan) |
| Zizyphi Fructus | Uchida Wakanyaku Co.,Ltd. (Tokyo, Japan) |
| Alismatis tuber | Tochimoto Tenkaido Co. Ltd. (Osaka, Japan) |
| Polyporus | Tochimoto Tenkaido Co. Ltd. (Osaka, Japan) |
| Ginseng Radix | Uchida Wakanyaku Co.,Ltd. (Tokyo, Japan) |
| Aconitii Radix Processa | Tochimoto Tenkaido Co. Ltd. (Osaka, Japan) |
| Poria | Tochimoto Tenkaido Co. Ltd. (Osaka, Japan) |

